# Lack of ancestral SARS-CoV-2 imprinting promotes BA.3.2.2 infection in children

**DOI:** 10.64898/2026.06.05.730251

**Authors:** Xiao Niu, Dan Xiong, Peizhuo He, Lingling Yu, Yutong Li, Xingan Cai, Ran An, Yao Wang, Ruoxi Kong, Yuanling Yu, Jing Wang, Qianran Wang, Binmao Zhang, Tianen Zhu, Tianyuan Zhao, Jianheng Huang, Fanchong Jian, Fei Shao, Zhimin Huang, Xiaosu Chen, Zhongyang Shen, Ronghua Jin, Xiaotian Tan, Yunlong Cao

## Abstract

The ongoing evolution of SARS-CoV-2 is heavily constrained by population-level immune imprinting. Recent genomic surveillance reveals an unexpected demographic shift: the highly mutated BA.3.2.2 sublineage is significantly enriched in paediatric populations globally, unlike concurrent variants XFG and NB.1.8.1. Here, the evidence demonstrates that this paediatric susceptibility arises from the absence of ancestral Wuhan-strain immune imprinting. Serological analyses reveal a profound failure to neutralise BA.3.2.2 in young children with no ancestral strain exposure. Concurrently, single-B-cell sequencing analyses demonstrate that BA.3.2.2 completely evades Omicron-specific Class 1/4 neutralising antibodies, encoded by IGHV2-5 and IGHV5-51, which dominate the repertoires of weakly imprinted individuals such as young children. Conversely, BA.3.2.2 remains uniquely susceptible to broadly cross-reactive Class 1 antibodies, encoded by IGHV3-53/66, typically enriched in adults with strong ancestral imprinting, such as mRNA vaccine recipients. Critically, sustained transmission of BA.3.2.2 in paediatric populations may catalyse the emergence of secondary variants that combine the paediatric-evading features of BA.3.2.2 with adult-evading Class 1 mutations. This could allow the lineage to breach the imprinted immunity of the adult population, potentially driving widespread global transmission. Together, these findings show that legacy immune imprinting paradoxically provides superior protection against a highly divergent saltation variant, directly explaining the age-biased transmission of BA.3.2.2, and informing future paediatric vaccination strategies.

## Main

The evolutionary landscape of Severe Acute Respiratory Syndrome Coronavirus 2 (SARS-CoV-2) has been characterised by continued adaptation of Omicron-derived lineages under widespread population immunity. During this period, the JN.1 descendant NB.1.8.1 and the recombinant lineage XFG became the major circulating variants since 2025 ^1–6^. The evolutionary success of both XFG and NB.1.8.1 is consistent with a broader pattern of incremental Omicron evolution, in which new variants acquire drifting mutations that reduce antibody neutralisation while maintaining optimal human angiotensin-converting enzyme 2 (hACE2) receptor binding ^2–4,7^.

In contrast, the BA.3.2 lineage and its descendants emerged as highly divergent saltation variants ^6,8,9^. First detected in Southern Africa in 2024 and later reported across multiple continents, BA.3.2 carries more than 50 spike mutations relative to BA.3 and more than 40 relative to JN.1, consistent with a saltation-like evolutionary pattern, potentially after prolonged intra-host evolution ^10^. Consistent with this, naïve animal immunization further revealed that BA.3.2.2 is antigenically distinct from JN.1-lineage viruses ^3,11^. Despite this antigenic divergence, BA.3.2.2 has remained less prevalent than XFG or NB.1.8.1, possibly reflecting fitness costs such as altered spike conformation, reduced entry efficiency, or slower replication ^3^. These features make BA.3.2.2 an unusual lineage with substantial antigenic distinction but limited overall spread.

However, the global prevalence of BA.3.2.2 has recently shown a marked upward trend since early 2026 (Fig. 1a) ^12^. More importantly, genomic surveillance identified a profound epidemiological anomaly: whereas XFG and NB.1.8.1 infections are concentrated in adult and elderly demographics, BA.3.2.2 infections exhibit a dramatic, statistically significant enrichment in paediatric populations (Fig. 1b, 1c and Extended Data Fig. 1a) ^13^. Generally, infectious respiratory diseases caused by SARS-CoV-2 variants heavily burden older populations owing to immunosenescence, cumulative comorbidities, and waning immunity ^14,15^. The inversion of this demographic vulnerability pattern suggests a highly specific immunological interaction between the unprecedented mutational profile of BA.3.2.2 and the distinct immunological histories of children.

**Figure 1.**
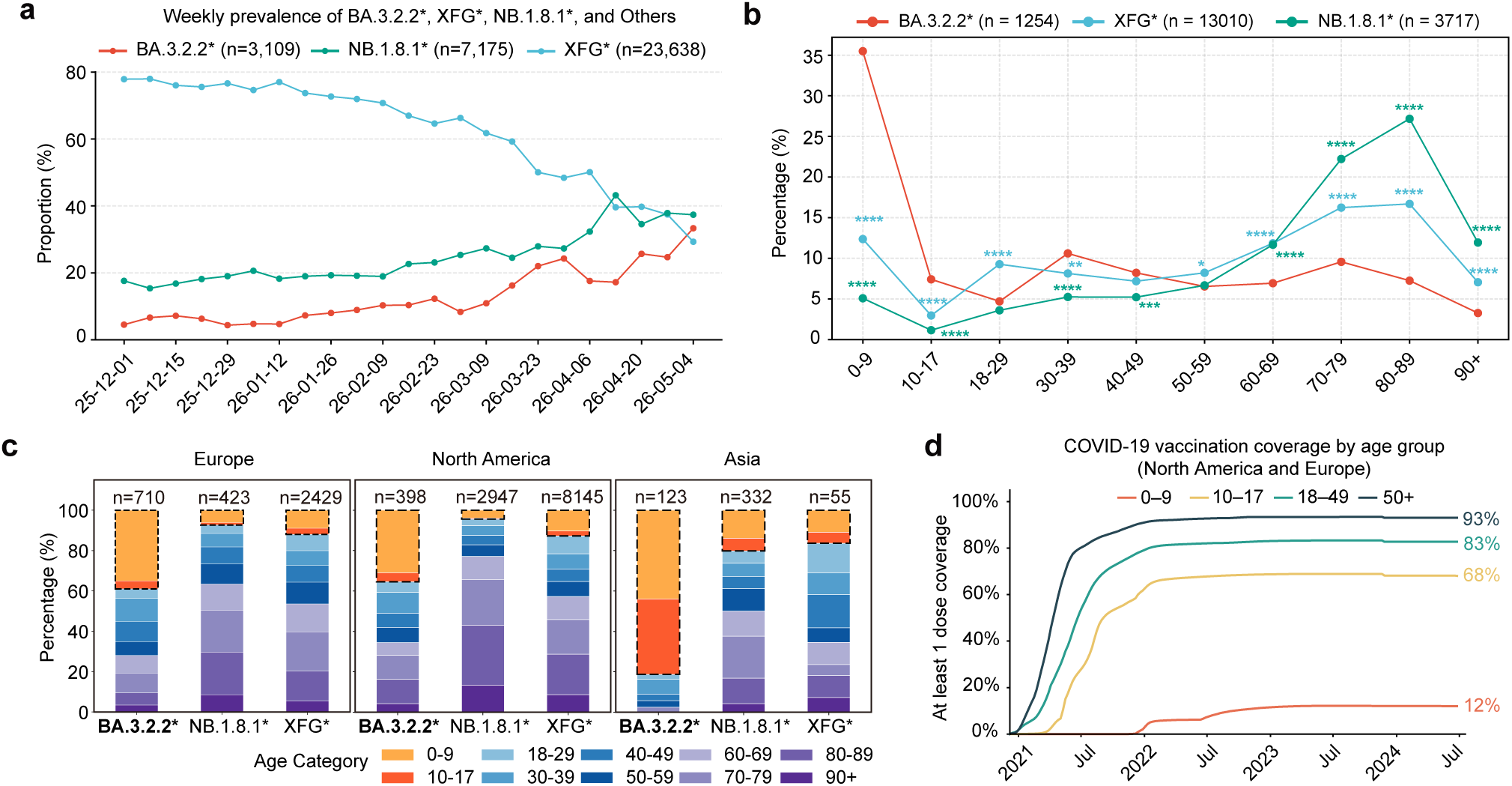
BA.3.2.2 is enriched in pediatric populations with low ancestral-strain exposure. **a**, Global weekly prevalence ratio of NB.1.8.1, XFG and BA.3.2.2 sublineages. **b**, Globally age distribution of NB.1.8.1, XFG and BA.3.2.2 sublineages. **c**, Bar plots showing the age distribution of BA.3.2.2, NB.1.8.1 and XFG sublineages globally and across continents. The group aged 17 years or younger is circled with black dashed outlines. Data in **a-c** were obtained from GISAID and include records collected between 1 December 2025 and 25 May 2026. **d**, Age-stratified vaccine coverage in major countries. Vaccination data were obtained from ECDC, the US CDC and canada.ca. Broadly defined age groups, including groups such as under 60 years and 18–69 years, were excluded. Chi-square tests were used in **b**.*P < 0.05, **P < 0.01, ***P < 0.001, ****P < 0.0001; ns, not significant (P > 0.05).

Immunological naivety provides an intuitive explanation for the apparent enrichment of BA.3.2.2 in younger populations. Children have generally had fewer SARS-CoV-2 exposures than adults, and might therefore be more susceptible to a lineage that is antigenically distant from previously circulating variants. However, a comparable age skew was not observed during the major antigenic transition from XBB-lineage viruses to BA.2.86/JN.1, despite the broad humoral immune escape and antigenic distinctiveness of the latter lineage. This contrast argues against immunological naivety alone as the primary driver of BA.3.2.2 enrichment in children.

From another perspective, fewer and later exposures to ancestral-strain vaccines in children also imply weaker immune imprinting (Fig. 1d and Extended Data Fig. 2a). In principle, weaker ancestral-strain imprinting would be expected to favour broader and more flexible neutralising responses to antigenically drifted Omicron lineages ^16–23^. The preferential enrichment of BA.3.2.2 in younger individuals, therefore, raises the possibility that this lineage exploits a more specific gap in the antibody landscape of individuals with limited ancestral-strain imprinting, whereas strongly imprinted adults may retain antibody specificities that better neutralise BA.3.2.2. This paradox prompted an investigation into whether imprinting-dependent differences in antibody repertoires underlie the age-associated spread of BA.3.2.2.

## Results

### Limited BA.3.2.2 neutralisation in children

To directly investigate the immunological basis of the paediatric enrichment of BA.3.2.2, two paediatric cohorts with distinct SARS-CoV-2 vaccination histories were recruited, and serum neutralisation against a panel of SARS-CoV-2 variant pseudoviruses was measured (Supplementary Table 1 and Fig. 2a). The first cohort consisted of unvaccinated children aged 8 years or younger (n = 16), representing the immune landscape of most young children, who had no exposure to the ancestral SARS-CoV-2 strain but had experienced natural Omicron infections. To serve as a comparison, the second cohort comprised children aged 8 to 14 years (n = 16) who had received priming with the inactivated Wuhan-strain vaccine. Given that China has recommended this primary vaccination for children older than 3 years since 2021, this cohort reflects the typical immune landscape of school-aged children.

**Figure 2.**
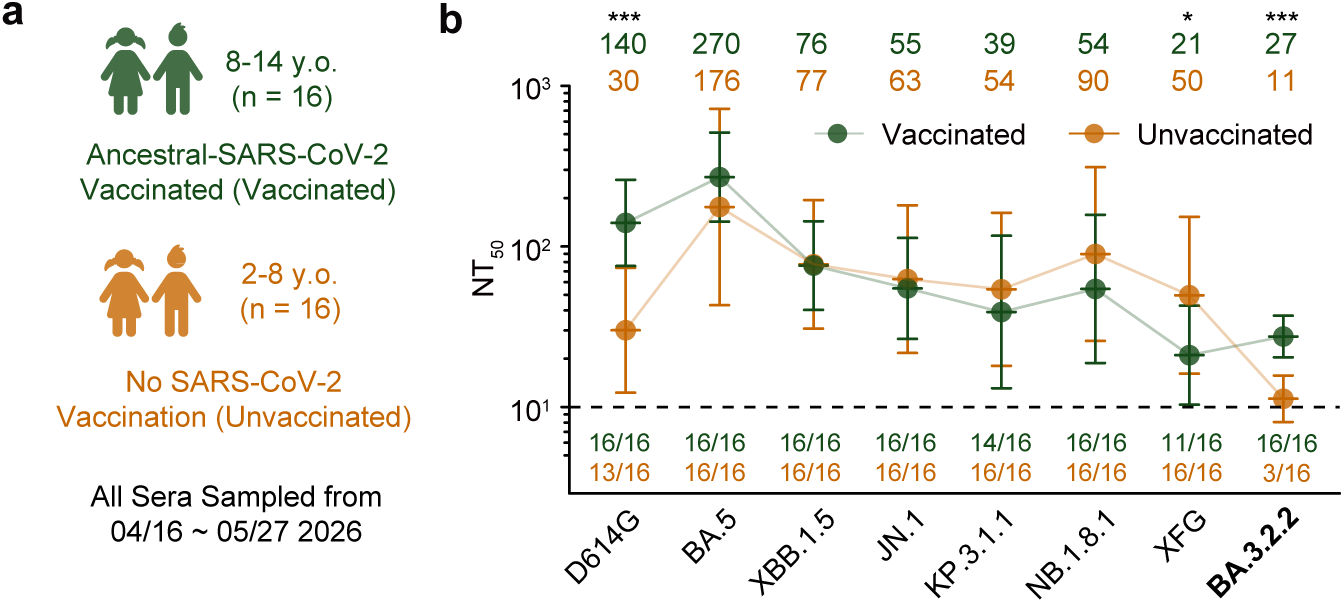
Limited BA.3.2.2 neutralisation in unvaccinated children. **a**, Schematic of the vaccinated and unvaccinated paediatric cohort included in this study. **b**, Comparison of neutralising antibody responses between the paediatric cohorts against a panel of SARS-CoV-2 pseudoviruses. Geometric mean titres (GMTs) are shown on the top. Dashed lines indicate the limit of detection (LOD, NT_50_ = 10). Data are presented as geometric mean titres (GMT), with error bars indicating geometric standard deviation. Positivity was defined as a titre above the limit of detection (NT_50_ = 10) and shown under the LOD. Two-tailed Wilcoxon rank-sum tests were used in **b**. *P < 0.05, **P < 0.01, ***P < 0.001, ****P < 0.0001.

Indeed, children aged 8 years or younger exhibited markedly less ancestral-strain SARS-CoV-2 immune imprinting than older children, as evidenced by significantly lower neutralising titres against D614G and overall stronger neutralisation against JN.1 sublineages, notably XFG (Fig. 2b). However, the unvaccinated children cohort demonstrated significantly lower neutralising titres against BA.3.2.2, with 13 of the 16 samples falling below the limit of detection. This diminished BA.3.2.2 neutralisation was not merely a reflection of globally weaker Omicron neutralisation; the vaccinated children cohort displayed robust neutralisation against BA.3.2.2, exceeding their titres against XFG. This neutralisation pattern aligns with the observed paediatric enrichment of BA.3.2.2 and raises the possibility that this variant preferentially exploits antibody repertoires shaped by weaker ancestral-strain immune imprinting.

### Imprinting severity dictates BA.3.2.2 neutralisation

If the severity of ancestral SARS-CoV-2 immune imprinting dictates BA.3.2.2 neutralisation, this phenomenon should be observable not only in unvaccinated children but also among adults with varying degrees of imprinting. Because mRNA vaccines induce strong imprinting while inactivated vaccines generate a more adaptable, Omicron-focused response following breakthrough infections ^24–32^, we examined a cohort of 20 adults with distinct vaccination histories. The 20 adults are long-term residents recruited from the same geographic region to reduce heterogeneity in local variant exposure (Supplementary Table 2 and Fig. 3a). The cohort was stratified by vaccination history: an inactivated-only group (n = 12; three CoronaVac doses) and an mRNA-vaccinated group (n = 8; ≥1 BNT162b2 or mRNA-1273 dose), with blood sampled approximately 46 months post-initial vaccination.

**Figure 3.**
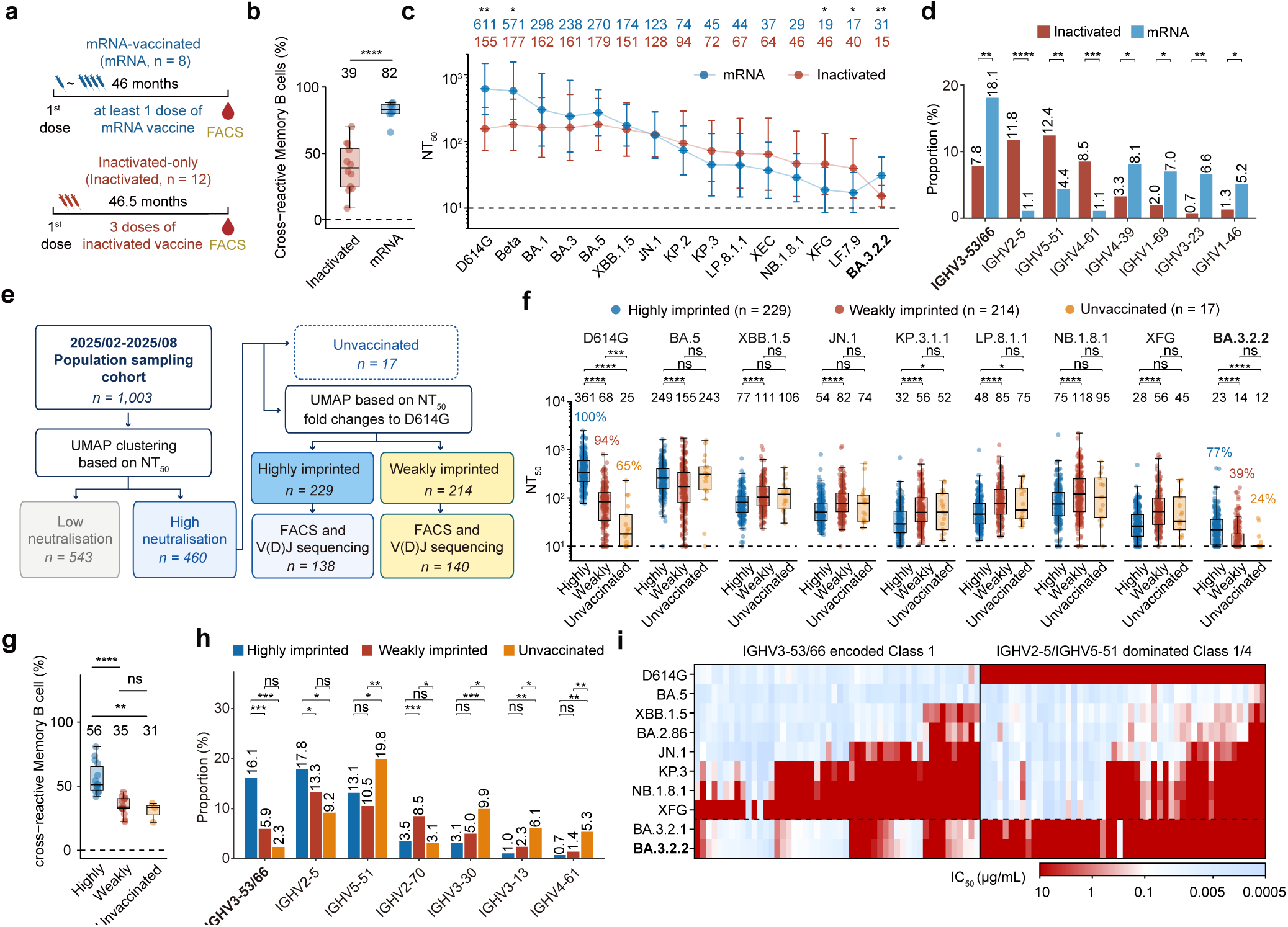
Ancestral-strain immune imprinting dictates BA.3.2.2 neutralisation. **a**, Immune histories and sampling timelines of the inactivated-only and mRNA-vaccinated cohorts. **b**, Proportion of Wuhan-cross-reactive (JN.1-positive) memory B cells. Boxes: median and IQR; whiskers: 1.5× IQR. **c**, Pseudovirus NAb responses for both cohorts. Top values indicate geometric mean titers (GMTs). Dashed lines: limit of detection (LOD, NT_50_=10). Error bars: geometric standard deviation. **d**, Heavy-chain V gene usage frequency (displaying significantly different genes representing >5% of sequences in both cohorts). **e**, Experimental and analysis workflow of randomly sampled human specimens. **f**, Pseudovirus NAb responses of the three groups. Top values indicate GMTs. Dashed lines: LOD. Plasma positivity rates (NT_50_ > 10) for D614G and BA.3.2.2 are noted. **g**, FACS analyses of the proportion of cross-reactive memory B cells (using a 10-in-1 sample pooling strategy). **h**, Heavy-chain V gene usage in highly imprinted, weakly imprinted, and unvaccinated cohorts (displaying significantly different genes representing >5% in cohorts). **i**, Heat map of variant escape profiles from Class 1 mAbs and Class 1/4 mAbs. Two-tailed Wilcoxon rank-sum tests were used in b-c and **f-g**. Chi-square tests were used in d and **h**. *P < 0.05, **P < 0.01, ***P < 0.001, ****P < 0.0001; ns, not significant (P > 0.05).

FACS analysis revealed that the mRNA-vaccinated group possessed a significantly higher frequency of Wuhan-cross-reactive memory B cells (MBCs) than the inactivated-only group (Extended Data Fig. 3a-3b and Fig. 3b). Serum Wuhan-RBD depletion also confirmed stronger ancestral-RBD cross-reactivity following mRNA vaccination, whereas Omicron-specific antibodies dominated the inactivated-vaccine response (Extended Data Fig. 3c). Pseudovirus neutralisation assays further highlighted distinct antibody profiles between the two groups (Fig. 3c). The mRNA group’s titres peaked against D614G and declined against subsequent variants, whereas the inactivated-only group displayed a flatter profile with significantly higher titres against recent variants like XFG ^33–37^. However, BA.3.2.2 displayed the inverse pattern. BA.3.2.2 was neutralised significantly more efficiently by sera from the mRNA-vaccinated group than by sera from the inactivated-only group (Fig. 3c). Thus, sera from both young children and weakly imprinted adults demonstrated selectively reduced BA.3.2.2 neutralisation, thereby linking the paediatric phenotype to imprinting-associated differences in humoral immunity.

This BA.3.2.2 selective neutralisation drive by imprinting level must stem from differences in the SARS-CoV-2 antigen-specific antibody repertoire landscape. To investigate this, we performed single-B-cell V(D)J sequencing of JN.1-reactive memory B cells. The strongly imprinted mRNA-vaccinated cohort was heavily enriched for IGHV3-53 and IGHV3-66 germline genes, which typically encode classical “public” Class 1 antibodies that target the ACE2-binding site ^38–45^ (Fig. 3d and Extended Data Fig. 4a). Such antibodies are readily elicited by ancestral-strain exposure but not Omicron infection, partly because their germline-encoded CDRH1 and CDRH2 motifs could naturally bind to the Wuhan Receptor-binding Domain (RBD). Conversely, the inactivated-only repertoire was dominated by IGHV5-51 and IGHV2-5 segments, which typify Omicron-specific Class 1/4 antibodies generated *de novo* upon antigenic shift ^16^. This suggests BA.3.2.2 preferentially escapes the Omicron-specific antibodies dominant in weakly imprinted individuals while remaining sensitive to ancestral-retained Class 1 antibodies.

Although the vaccination-history cohorts identified imprinting-associated differences in BA.3.2.2 neutralisation and BCR gene usage, the analysis was limited by small cohort size. To validate these imprinting-associated phenotypes in a broader population, we analysed 1,003 randomly collected blood samples (February–August 2025, Beijing) for downstream pseudovirus neutralisation assays (Supplementary Table 2 and Fig. 3e). Overall, NT_50_ values against all tested SARS-CoV-2 variants increased over the sampling period in this randomly sampled cohort (Extended Data Fig. 5a-5b), likely reflecting the accumulation of NB.1.8.1 exposures in the population. To minimise the influence of low-neutralisation samples on later fold-change analyses and statistical comparisons, we first performed UMAP clustering using NT_50_ values against variants to separate the cohort into high- and low-neutralisation clusters (Extended Data Fig. 6a). Among the high-neutralisation samples, we excluded 17 unvaccinated individuals and then performed a second UMAP analysis based on NT_50_ fold changes of the eight variants relative to D614G. This further classified the remaining 443 samples into highly imprinted (D614G-biased) and weakly imprinted groups (Omicron-specific) (Fig. 3f and Extended Data Fig. 6b). FACS analyses of memory B cell cross-reactivity confirmed the validity of this grouping (using a 10-in-1 sample pooling strategy), as the highly imprinted group displayed a significant enrichment of Wuhan-reactive, JN.1-specific memory B cells (Fig. 3g).

Consistent with the smaller cohorts, BA.3.2.2 exhibited stronger neutralisation escape in both the weakly imprinted and unvaccinated groups compared to the highly imprinted group (Fig. 3f). Furthermore, unvaccinated individuals showed robust neutralisation of most Omicron variants but significantly lower positivity against BA.3.2.2, firmly linking weak Wuhan-strain imprinting to BA.3.2.2 escape (Extended Data Fig. 7a, b). Subsequent V(D)J sequencing of JN.1-specific memory B cells confirmed that highly imprinted individuals again possessed elevated IGHV3-53/66 usage, whereas these gene segments were largely absent in unvaccinated individuals (Fig. 3h and Extended Data Fig. 8a)

To definitively link these distinct antibody classes distinguished by IGHV usage to the BA.3.2.2 escape phenotype, we evaluated the neutralisation profile of a panel of 100 recombinant monoclonal antibodies (mAbs) comprising 50 IGHV3-53/66-encoded Class 1 mAbs and 50 IGHV2-5/IGHV5-51-dominated Class 1/4 mAbs. As predicted, the IGHV3-53/66-encoded Class 1 mAbs (enriched in highly imprinted individuals) broadly retained neutralisation against BA.3.2.1 and BA.3.2.2. In stark contrast, the Class 1/4 mAbs (enriched in weakly imprinted individuals) were frequently escaped, with only two retaining activity against BA.3.2.2 (Fig. 3i).

Collectively, these data provide a mechanistic explanation for the age-biased distribution of BA.3.2.2. By escaping Omicron-specific Class 1/4 antibodies while remaining sensitive to IGHV3-53 encoded Wuhan-cross-reactive Class 1 antibodies, BA.3.2.2 exploits the weakly imprinted immune landscapes typical of young children (who have lower or no ancestral-strain vaccine coverage). Furthermore, while limited sequence deposition and sustained NB.1.8.1 predominance may explain the current lack of widespread BA.3.2.2 detection in China, the population’s increasing prevalence of weakly imprinted immune histories may create a permissive landscape for future BA.3.2.2 expansion.

### Potential BA.3.2.2 evolution

The increasing prevalence of BA.3.2.2 in the paediatric population could create a large reservoir for viral replication and accelerated evolution. Similar to the evolutionary trajectory from BA.2.86 to JN.1, this could eventually enable future BA.3.2.2 sublineages to efficiently infect adult and elderly populations. To assess whether BA.3.2.2 could further expand its immune evasion after entering these weakly imprinted populations, we generated a panel of BA.3.2.2 pseudoviruses carrying additional RBD substitutions. This panel included mutations observed in contemporary Class 1 antibody-evading Omicron lineages and a Y505H reversion mutant designed to test if this specific residue drives Class 1/4 antibody escape.

Evaluating these mutants against our monoclonal antibody (mAb) panel revealed that most substantially escaped Class 1 antibodies, especially the 455-456 “FLip” mutation combo (Fig. 4a) ^46,47^. Notably, the Y505H reversion restored neutralisation for the majority of Class 1/4 mAbs that parental BA.3.2.2 had completely escaped (Fig. 4b). These mAb results aligned perfectly with our plasma assays. Plasma from the highly imprinted group was significantly evaded by nearly all BA.3.2.2 mutants—often reaching escape levels comparable to XFG—indicating that additional RBD substitutions readily erode their residual neutralising immunity (Fig. 4c). Conversely, the Y505H reversion increased plasma neutralisation in the weakly imprinted and unvaccinated groups, but not in the highly imprinted group, confirming that the Y505H substitution is the key determinant of BA.3.2.2 escape from Omicron-specific antibody repertoires.

**Figure 4.**
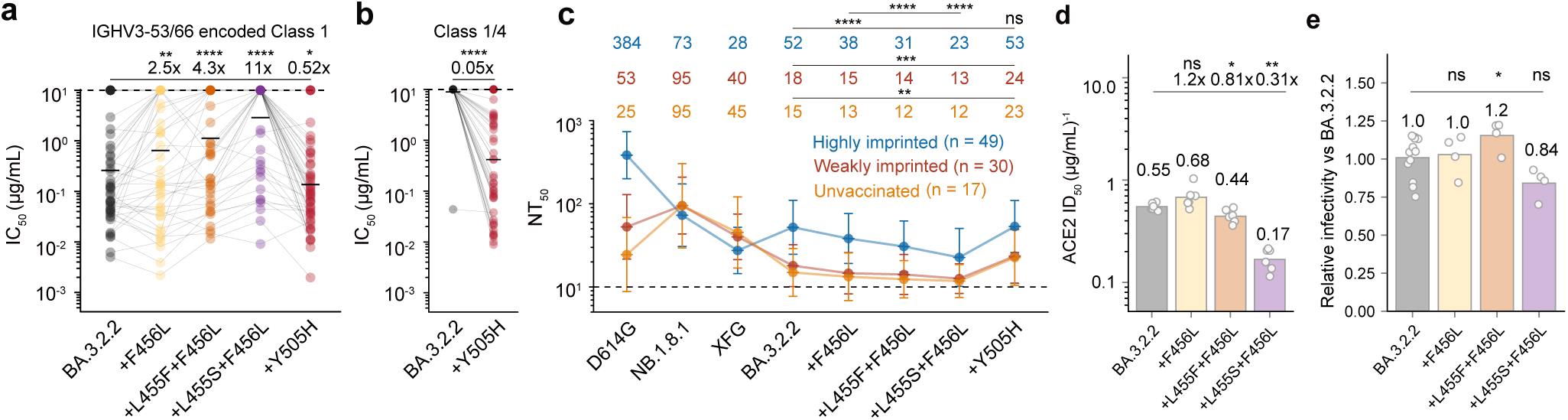
RBD substitutions enable further escape of BA.3.2.2 from strongly imprinted immunity. **a**, Scatter-line plot showing escape profiles of different BA.3.2.2 mutants from Class 1 mAbs encoded by IGHV3-53/66. **b**, Neutralisation recovery of Class 1/4 mAbs dominated by IGHV2-5 and IGHV5-51 by Y505H reverse mutation. **c**, NAb response of the randomly selected serum from BA.3.2.2-positive highly imprinted, weakly imprinted and unvaccinated individuals against a panel of BA.3.2.2 mutants pseudoviruses. Dashed lines indicate the limit of detection (NT_50_ = 10). Data are presented as geometric mean titers (GMT), with error bars indicating geometric standard deviation. Points represent individual samples.49 BA.3.2.2-neutralising plasma samples from the highly imprinted group, 30 samples from the weakly imprinted group, and all 17 samples from the unvaccinated group are randomly selected. **d**, ACE2 ID50 against a panel of BA.3.2.2 mutants pseudoviruses. **e**, Relative infectivity of BA.3.2.2 mutants in Vero cells. Infectivity was assessed by use of vesicular stomatitis virus pseudoviruses. Two-tailed Wilcoxon rank-sum tests were used in **a-e**. *P < 0.05, **P < 0.01, ***P < 0.001, ****P < 0.0001; ns, not significant (P > 0.05).

To determine if these escape routes compromise viral fitness, we quantified receptor engagement by measuring ACE2 inhibition of these BA.3.2.2 mutant pseudoviruses and assessed spike-mediated entry using a VSV-based pseudovirus infectivity assay in Vero cells. While some substitutions (e.g., L455S) heavily impaired receptor engagement, F456L—alone or combined with L455F—maintained or slightly enhanced ACE2 binding and entry efficiency relative to parental BA.3.2.2 (Fig. 4d, 4e). Thus, F456L-related changes provide a plausible route by which BA.3.2.2 could acquire broader antibody escape while retaining, or potentially improving, receptor engagement and entry efficiency ^48^.

## Discussion

Together, our findings reveal an unusual relationship between immune imprinting, antibody-repertoire composition, and the age-biased distribution of BA.3.2.2. Whereas strong ancestral-strain imprinting generally constrains adaptation to recent Omicron variants (e.g., NB.1.8.1 and XFG), BA.3.2.2 exhibits the opposite pattern. It preferentially escapes the Omicron-specific Class 1/4 antibodies enriched in weakly imprinted individuals, while remaining sensitive to the IGHV3-53/66-encoded, Wuhan-cross-reactive Class 1 antibodies enriched in strongly imprinted individuals. This antibody-class specificity mechanistically explains the paediatric enrichment of BA.3.2.2, given that children generally have lower ancestral-strain vaccine coverage and correspondingly weaker Wuhan-strain imprinting.

A critical insight derived from these observations is that legacy immune imprinting does not act as a uniform barrier to neutralising efficacy; rather, its utility is highly dependent on the precise conformation of the emerging variant’s antigenic landscape. While imprinting has historically been characterised as a constraint that leaves populations vulnerable to incremental drift, it serves here as a robust protective shield against specific saltation variants. This is due to the broad-spectrum neutralisation of germline-encoded Class 1 public antibodies, which the virus has yet to successfully bypass. In contrast, the *de novo*, Omicron-specific responses that dominate the low-imprinted populations are highly specialised but narrow, rendering them vulnerable to BA.3.2.2.

This points to a significant public health vulnerability. Paediatric cohorts, lacking ancestral-strain exposures, provide an immunological niche—an incubator—where BA.3.2.2 can propagate with minimal resistance. Within this weakly imprinted population, the genetic barrier for the virus to acquire supplementary mutations (such as F456L-related changes) is extremely low. This raises a concern: sustained transmission of BA.3.2.2 in paediatric reservoirs may catalyse the emergence of secondary variants that combine the paediatric-evading features of BA.3.2.2 with adult-evading Class 1 mutations. Such evolutionary trajectories would allow the lineage to breach the imprinted immunity of the adult population, potentially driving widespread global transmission.

Thus, the impact of immune imprinting depends heavily on how viral antigenic change intersects with stratified population-level antibody repertoires. These findings underscore that future paediatric SARS-CoV-2 vaccination campaigns should weigh the immediate benefits of variant-matched antigens against the long-term utility of establishing a broad antibody baseline. There is a critical need for immunization strategies that combine variant-matched specificity with legacy-like breadth to ensure balanced protection across all age groups.

## Data availability

Information on individuals providing blood samples and neutralisation titres involved in this study has been included in the Supplementary Table. Further information and requests for resources and reagents should be directed to and will be fulfilled by the lead contact, (Y.C.).

## Code availability

No custom code was developed for this study. Data analysis was performed using standard software or packages/libraries in R or python, as detailed in the Methods section.

## Acknowledgments

We thank all volunteers who provided blood samples. This project is financially supported by Chinese Academy of Sciences (to X.T.) and Changping Laboratory (2026D-04-01 to Y.C.).

## Author contributions

Y.C. designed and supervised the study. X.N. and Y.C. wrote the manuscript with input from all authors. X.N. and P.H. curated and analysed the epidemiological data. X.N., R.A. and Y.W. performed B cell sorting, 10x single-cell V(D)J sequencing experiments and data analysis. Q.W. performed plate-based single B cell V(D)J sequencing. X.N., Y.L., X.C., R.A., Y.W., and R.K. performed FACS analysis. J.W. and F.S. performed antibody expression. Y.Y. and P.H. constructed the pseudotyped virus and performed the pseudovirus infectivity assays. L.Y. performed the pseudovirus neutralisation assays and ELISAs. F.J., P.H. and X.N. constructed the BA.3.2.2 mutant plasmids. D.X., Z.H., T.Z., and X.T. recruited the children cohort and collected the blood samples. B.Z., T.Z., J.H., and X.T. recruited the mRNA vaccinated patients and collected the blood samples. X.C., Z.S. and R.J. recruited the randomly collected cohort and collected the blood samples.

## Methods

### Human serum and PBMC isolation

Blood samples were obtained from convalescent individuals who had received various vaccine platforms (detailed in Supplementary Table 1). The research protocol and the collection of human blood samples were approved by the Institutional Review Board (IRB) of Shenzhen Institutes of Advanced Sciences, Chinese Academy of Science (Ethics committee archiving no. SIAT-IRB-230715-H0667), the Institutional Review Board of Shenzhen Luohu People’s Hospital (Ethics committee archiving no. 2260206006), the Tianjin Municipal Health Commission and the Ethics Committee of Tianjin First Central Hospital (Ethics committee archiving no. KEYAN20241022-2), and Beijing Ditan Hospital, Capital Medical University (Ethics committee archiving no. DTEC-KY2024-112-01). All participants provided their agreement for the collection, storage and use of their blood samples strictly for research purposes and the subsequent publication of related data.

To isolate serum and PBMCs, whole blood was diluted 1:1 with PBS (Invitrogen, C10010500BT) containing 2% (v/v) fetal bovine serum (FBS; Hyclone, SH30406.05) and separated by density gradient centrifugation using Ficoll (Cytiva, 17-1440-03). The upper serum layer was collected, aliquoted, and stored at or below −20°C. Before use in assays, serum samples were heat-inactivated at 56°C for 30 min and assessed for neutralising titres against SARS-CoV-2 variant spike-pseudotyped vesicular stomatitis virus (VSV).

The PBMC layer was harvested from the interface and processed further. Following red blood cell lysis (Invitrogen eBioscience 1X RBC Lysis Buffer, 00-4333-57) and washing, PBMCs were either used immediately or cryopreserved in FBS with 10% (v/v) DMSO (Solarbio, D8371) for storage in liquid nitrogen. All PBMC samples were transported on dry ice. For B cell isolation, cryopreserved PBMCs were thawed in PBS supplemented with 5% (v/v) FBS. B cells were subsequently enriched by immunomagnetic positive selection using the EasySep™ Human CD19 Positive Selection Kit II (STEMCELL, 17854) following the manufacturer’s protocol. The resulting purified B cells were resuspended in PBS with 2% (v/v) FBS, and cell counts and viability were determined using 0.4% (w/v) trypan blue staining (Invitrogen, T10282) on a Countess Automated Cell Counter (Invitrogen).

### Pseudovirus preparation and neutralisation assay

We generated SARS-CoV-2 variant spike protein pseudovirus as described previously ^3,16,18,49–51^. Plasmids encoding a codon-optimised SARS-CoV-2 Spike (S) protein were constructed by inserting the S gene into the pcDNA3.1 vector. To produce pseudovirus, 293T cells (ATCC, CRL-3216) were transfected with the Spike-expressing plasmids with Lipofectamine 3000 (Invitrogen, L3000015) and subsequently infected with G*ΔG-VSV (Kerafast, EH1020-PM). After 24 hours, the supernatant containing the pseudovirus was harvested, filtered through a 0.45 μm filter (Millipore), aliquoted, and stored at −80°C.

Neutralisation assays were performed using the Huh-7 cell line (JCRB, 0403). Monoclonal antibodies or serum samples were serially diluted in DMEM (Hyclone, SH30243.01) and incubated with the pseudovirus in 96-well plates for 1 hour at 37°C with 5% CO₂. Following incubation, Huh-7 cells were seeded into the wells (2×10^4^ cells per well) and cultured for an additional 24 hours at 37°C with 5% CO₂. To assess infection levels, the culture supernatant was removed and left 50 μl in each well. The Bright-Lite Luciferase Assay Substrate was reconstituted with its corresponding Assay Buffer (Vazyme, DD1209-03-AB), and this mixture was added to the wells. After incubating in the dark for 2 minutes, luminescence was measured using a microplate spectrophotometer (PerkinElmer, HH3400). The NT_50_ or IC_50_ values were determined using a three-parameter logistic regression model.

### Flow cytometry analysis and antigen-specific B cell sorting

For the isolation of antigen-specific human B cells, enriched B cell populations from PBMCs were prepared for fluorescence-activated cell sorting (FACS). The staining panel included FITC anti-human CD20 (BioLegend, 302304), Brilliant Violet 605™ anti-human CD27 (BioLegend, 302824), PE/Cyanine7 anti-human IgM (BioLegend, 314532), and PE/Cyanine7 anti-human IgD (BioLegend, 348210). Antigen-specific cells were detected using biotinylated JN.1 RBD (Sino Biological, 40592-V49H16-B) conjugated with PE-streptavidin (BioLegend, 405204) and APC-streptavidin (BioLegend, 405207), and Wuhan RBD (Sino Biological, 40592-V27H-B) conjugated with BV421-streptavidin (BioLegend, 405225). The viability dye 7-AAD (Invitrogen, 00-6993-50) was used to exclude dead cells. A gating strategy was applied to sort single, viable (7-AAD⁻), class-switched (IgM⁻ and IgD⁻), CD20⁺CD27⁺ memory B cells that were positive for JN.1 RBD. Data were collected via Summit 6.0 software (Beckman Coulter). Data from all experiments were uniformly analysed using FlowJo v10.8 (BD Biosciences).

### Single-cell V(D)J sequencing

For the 10x Genomics workflow, sorted antigen-specific B cells, suspended in PBS with 10% (v/v) FBS, were processed with the Chromium Next GEM Single Cell V(D)J Reagent Kits v2 (10X Genomics, CG000331). The cell suspension was loaded onto a 10X Chromium Controller to generate Gel Beads-in-Emulsion (GEMs), which facilitate the barcoding of mRNA and subsequent reverse transcription within individual droplets. Following cDNA synthesis, the product was purified using a SPRIselect Reagent Kit (Beckman Coulter, B23318) and pre-amplified. Targeted enrichment of paired V(D)J sequences was then achieved using 10X-specific BCR primers, and the resulting products were used for sequencing library construction. These final libraries were sequenced on an Illumina NovaSeq 6000 platform with a NovaSeq 6000 S4 Reagent Kit v1.5 (300 cycles; Illumina, 20028312). The 10X Genomics V(D)J Illumina sequencing data were assembled as B cell receptor contigs and aligned to the B cell V(D)J reference using Cell Ranger (v.6.1.1) pipeline. For human-source IGH, IGL and IGK contigs, we used GRCh38 as reference. Only the productive contigs and B cells with one heavy chain and one light chain were kept to remove doublets. The germline V(D)J genes were identified and annotated using IgBlast (v1.17.1) ^52^.

For the plate-based method, single antigen-specific B cells were sorted directly into the individual wells of 96-well plates, each containing 4 µL of lysis buffer (0.095% Triton X-100, 2.5 µM oligo-dT primer: 5’-AAGCAGTGGTATCAACGCAGAGTACTTTTTTTTTTTTTTTTTTTTTTTTTTTTTTVN-3’, 2.5 mM dNTPs, 1 U/µL RNase Inhibitor). The plates were immediately processed by vortexing, centrifugation, and incubation at 72°C for 3 min to denature RNA secondary structures, followed by rapid chilling on ice. Reverse transcription was performed in each well by adding 5.7 µL of an RT master mix (final concentrations: 1× HiScript III first-strand buffer, 10 U/µL HiScript III reverse transcriptase, 2 U/µL RNase inhibitor, 5 mM DTT, 1 M betaine, 6 mM MgCl₂, 1 µM TSO primer: 5’-AAGCAGTGGTATCAACGCAGAGTACATrGrG+G-3’). The RT reaction was incubated at 37°C for 60 minutes, followed by enzyme inactivation at 85°C for 5 seconds. The resulting cDNA served as a template for amplifying full-length immunoglobulin heavy and light chain V(D)J sequences via a two-round nested PCR strategy using Phanta Max Master Mix (Vazyme, P515). The first PCR round employed a multiplex primer pool targeting all VH, Vκ, and Vλ gene families, along with constant region-specific reverse primers. A second, nested PCR round was then performed with internal primers to enhance specificity and yield. Final amplicons were purified and subsequently analysed by Sanger sequencing. The germline V(D)J genes were identified and annotated using IgBlast (v1.22.0) ^52^.

### Monoclonal antibody expression and purification

The sequences for the antibody heavy and light chains were initially codon-optimised for expression in human cells (GenScript). The variable regions (VH and VL) were then separately inserted into corresponding expression vectors (pCMV3-CH, pCMV3-CL or pCMV3-CK). Plasmids of the heavy and light chain constructs were transformed into *Escherichia coli* DH5α competent cells (Tsingke, TSC-C01-96). After overnight incubation at 37°C, single colonies were picked for colony PCR identification. Plasmid DNA of expanded cultures was extracted (CWBIO, CW2105) being after verified by Sanger sequencing.

For protein production, heavy and light chain plasmids were co-transfected into Expi293F cells (Thermo Fisher, A14527) using polyethylenimine (PEI; Yeasen, 40816ES03). The plasmid-PEI complexes were prepared in sterile 0.9% NaCl solution before being added to the cell culture. The transfected cells were cultured at 36.5°C with 5% CO₂ and 175 rpm shaking for 6–10 days. A nutrient supplement (OPM Biosciences, F081918-001) was added to each culture at 24 hours post-transfection and every 48 hours thereafter.

To purify the antibodies, the culture supernatant was first clarified by centrifugation (3,000 × g, 10 minutes). The supernatant was then incubated with Protein A magnetic beads (GenScript, L00695) for 2 hours to allow antibody binding. The beads were subsequently washed, and the bound antibodies were eluted using a KingFisher automated purification system (Thermo Fisher). The concentration of the purified antibody was determined using a NanoDrop spectrophotometer (Thermo Fisher, 840-317400), and its purity was assessed by SDS-PAGE (LabLead, P42015).

### RBD depletion of serum

To deplete RBD-specific antibodies from serum, 50 μL of Dynabeads™ MyOne™ Streptavidin T1 (Invitrogen, 65601) were washed once with PBS. 10 μg of biotinylated SARS-CoV-2 Wuhan RBD was incubated with the washed beads for 1 hour with gentle rotation to allow binding via streptavidin-biotin interaction. The beads were then collected using a magnetic rack for 2-3 min, the supernatant was discarded, and the beads were washed three times with PBS to remove unbound proteins. Subsequently, 200-400 μL of serum was incubated with the RBD-conjugated beads for 1 hour with gentle rotation to allow specific antibody binding. Finally, the tubes were placed on the magnetic rack, and the supernatant representing the RBD-depleted serum was carefully collected for downstream analyses.

### Enzyme-linked immunosorbent assays

High-binding 96-well plates (NEST, 504201) were coated overnight at 4°C with SARS-CoV-2 Wuhan (Sino Biological, 40592-V27H-B) or JN.1 RBD proteins (Sino Biological, 40592-V49H16-B). The following day, plates were washed three times with 1×PBST (Solarbio, P1033) and blocked with 250 μL 5% bovine serum albumin (BSA; Solarbio, A8020) in 1×PBST for 2 hours at 37°C to prevent non-specific binding. After three additional washes, 100 µL of serially diluted antibodies or serum samples were added to the wells and incubated for 30 minutes at room temperature. Unbound antibodies were removed by five washes with 1×PBST. Subsequently, 100 µL of HRP-conjugated Goat anti-Mouse IgG (H+L) Cross-Adsorbed Secondary Antibody (Invitrogen, G21040) or Peroxidase AffiniPure Goat Anti-Human IgG (H+L) (Jackson Immunoresearch, 109-035-003) was added and incubated for 30 minutes at room temperature. Following a final five washes, the signal was developed by adding 100 µL of TMB substrate (Solarbio, PR1200) to each well and incubating for 8 minutes in the dark. The reaction was terminated by adding 50 µL of stop solution (Solarbio, C1058). The optical density (OD) was measured at 450 nm with a reference wavelength of 630 nm using a Multiskan FC microplate reader (Thermo Scientific). Final absorbance values were obtained by subtracting the OD630 reading from the OD450 reading for each well.

### GISAID sequence retrieval and metadata curation

SARS-CoV-2 sequence metadata were obtained from the GISAID EpiCoV database. All available records with collection dates from 1 December 2025 to 25 May 2026 were screened, and records assigned to XFG*, BA.3.2.2* or NB.1.8.1* were retained for analysis. The asterisk denotes each parental Pango lineage together with all descendant lineages. Pango lineage annotations available in the GISAID metadata were used for initial lineage classification. The lineage_notes.txt and alias_key.json files obtained from the cov-lineages/pango-designation GitHub repository were subsequently used to expand aliased Pango lineages and identify all descendant lineages of the target parental lineages. Metadata were curated by excluding records with missing or non-informative collection date, sampling location, patient age or lineage annotations. Patient age entries were standardised to integer years where possible; infant ages reported in days, weeks or months were converted to 0 years, and uninterpretable or implausible values were removed. Harmonised ages were grouped into age bands from 0–4 to 80–89 years, with individuals aged ≥90 years assigned to a single upper age category.

### Age-stratified vaccination coverage data

Age-stratified COVID-19 vaccination coverage data were obtained from official public health surveillance sources, including the European Centre for Disease Prevention and Control for EU/EEA countries, the Government of Canada COVID-19 vaccination coverage database for Canada and the US Centers for Disease Control and Prevention vaccination surveillance dataset for the United States. These datasets were used to reconstruct retrospective age-stratified vaccine coverage during the period of primary ancestral-strain vaccination campaigns. For each region or country, vaccination coverage was extracted by reported age group and time point. Where necessary, age groups were harmonised to enable comparison with the age categories used in the sequence-based analyses. Broadly defined age groups that could not be assigned to comparable age strata, including categories such as under 60 years or 18–69 years, were excluded. Vaccine coverage was analysed descriptively to compare the timing and magnitude of vaccine uptake between children, adolescents, adults and older adults.

---

**Extended Data Figure 1.**
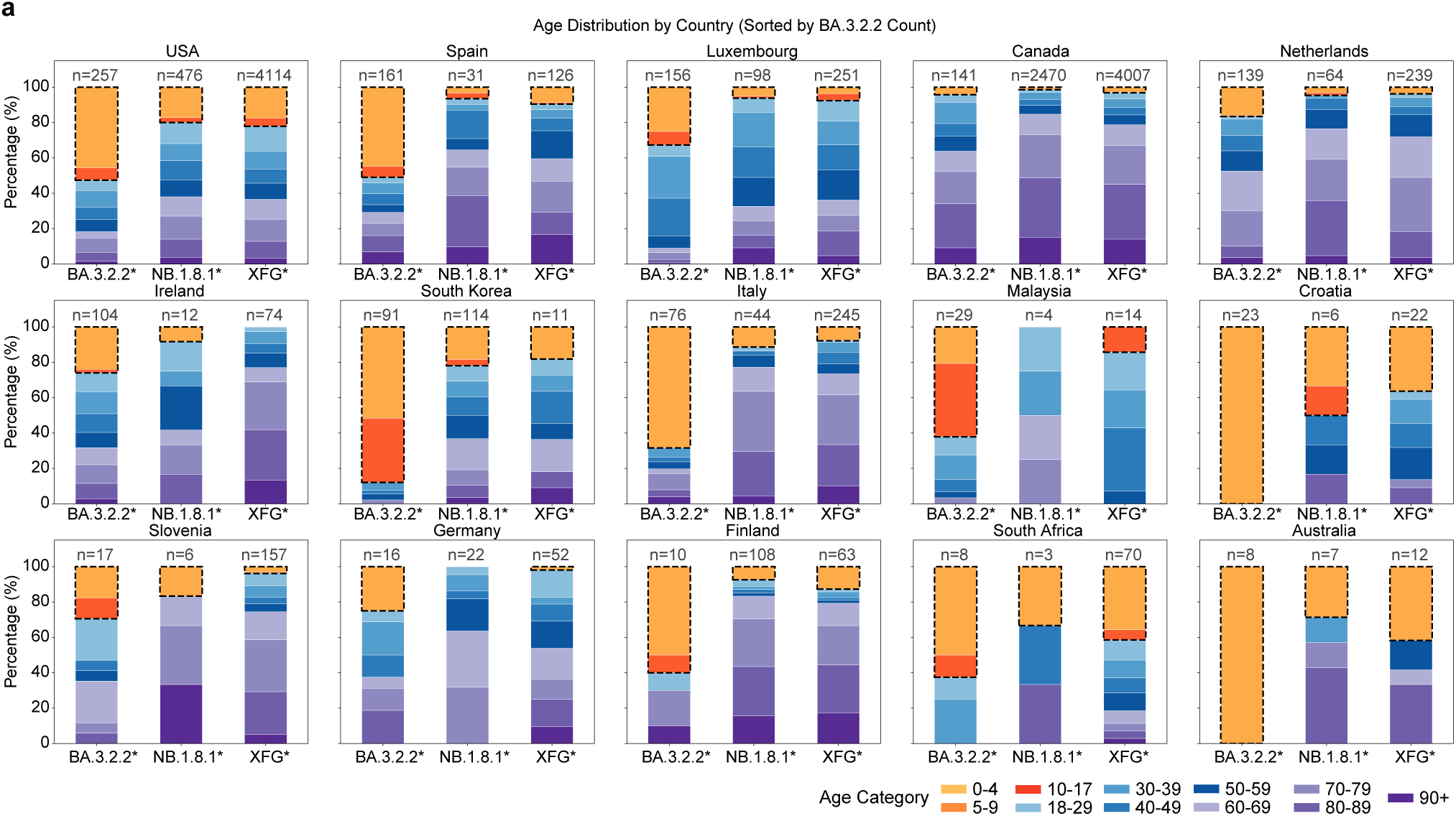
Country-level surveillance confirms the paediatric enrichment of BA.3.2.2. **a**, Bar plots showing the age distribution of BA.3.2.2, NB.1.8.1 and XFG sublineages across countries. The group aged 17 years or younger is circled with black dashed outlines. Data were obtained from GISAID and include sequences collected between 1 December 2025 and 25 May 2026. Data processing is described in the Methods.

**Extended Data Figure 2.**
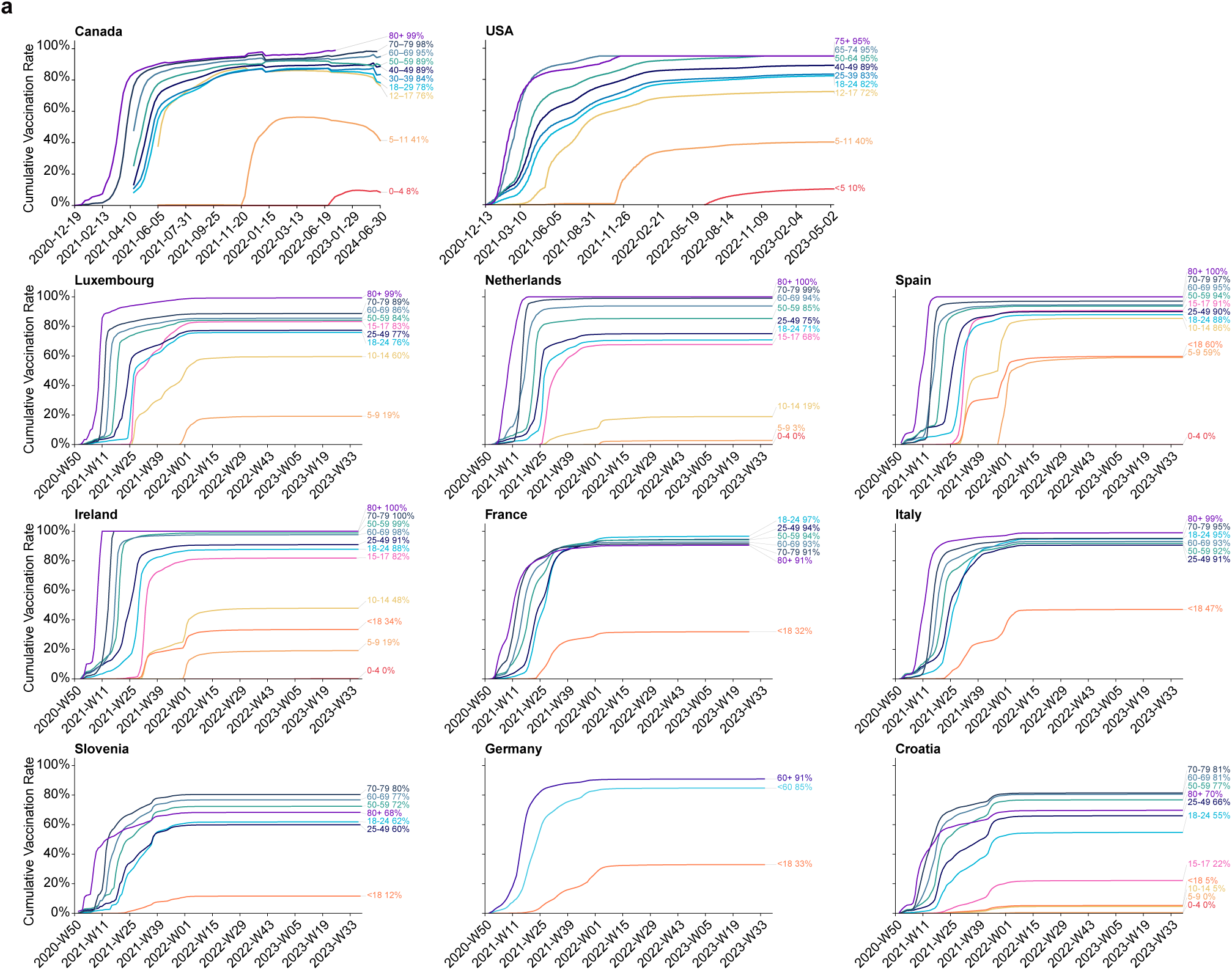
Children show consistently lower vaccine coverage across countries. **a**, Temporal changes in vaccine coverage across age groups in major countries. Vaccination data were obtained from ECDC, the US CDC and canada.ca. Broadly defined age groups, including groups such as under 60 years and 18–69 years, were excluded.

**Extended Data Figure 3.**
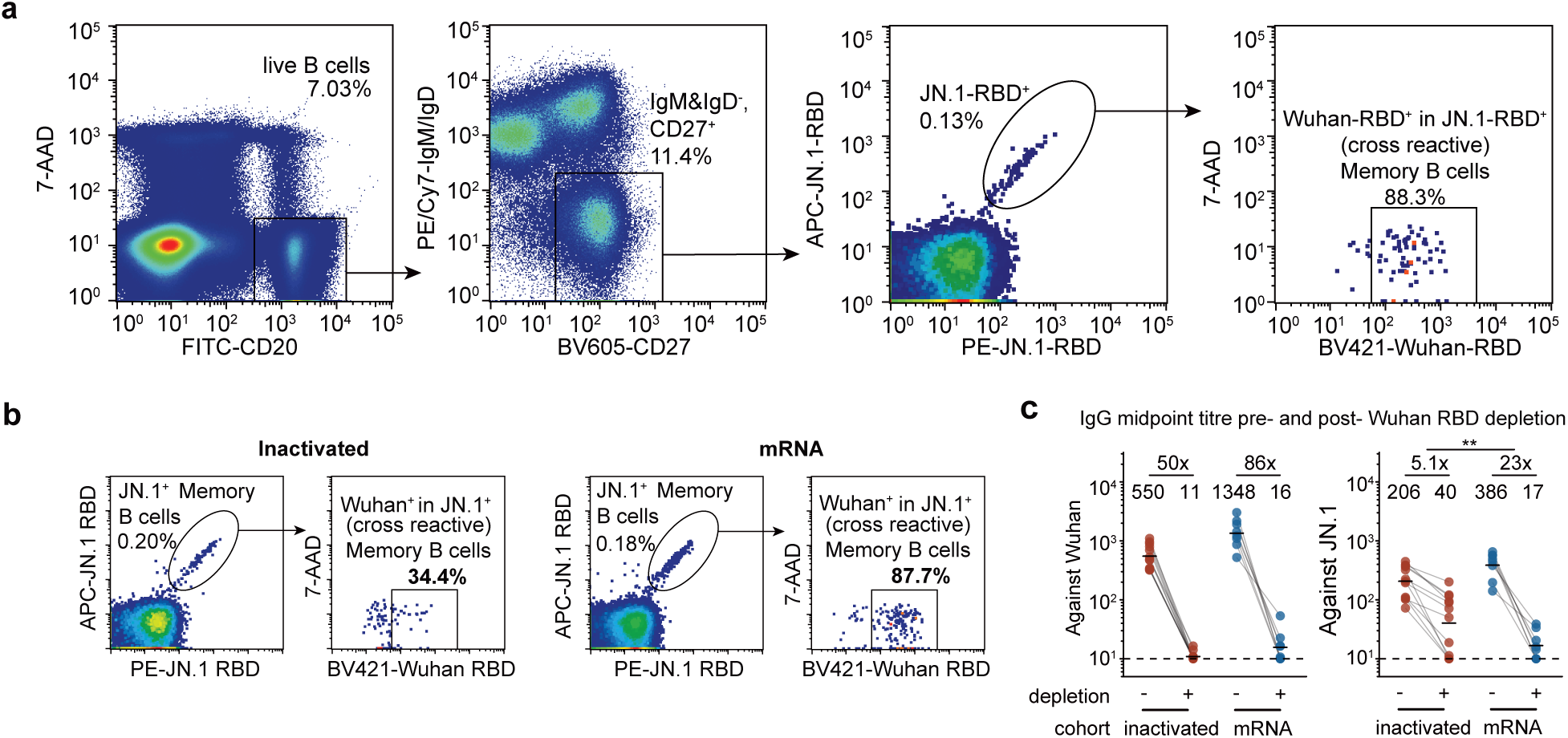
Cellular and serological profiling validates stronger ancestral imprinting in mRNA-vaccinated individuals. **a**, Representative gating strategy for sorting human B cells. FITC, fluorescein isothiocyanate; BV605, Brilliant Violet 605; PE/Cy7, PE/Cyanine7. **b,** Representative flow cytometry dot plots of cross-reactive memory B cells from the inactivated-only (top) and mRNA-vaccinated (bottom) cohorts. APC, allophycocyanin; PE, phycoerythrin; BV421, Brilliant Violet 421. **c**, Serum IgG midpoint titre against Wuhan (left) or JN.1 (right) RBD before and after Wuhan RBD depletion. Geometric mean values are displayed as bars and indicated above each group of data points. Statistical significance of the fold-reduction in titres was assessed between two cohorts. Dashed lines indicate the limit of detection (midpoint titre = 10). Two-tailed Wilcoxon rank-sum tests were used in **c**. \**P* < 0.05, \*\**P* < 0.01, \*\*\**P* < 0.001, \*\*\*\**P* < 0.0001; ns, not significant (P > 0.05).

**Extended Data Figure 4.**
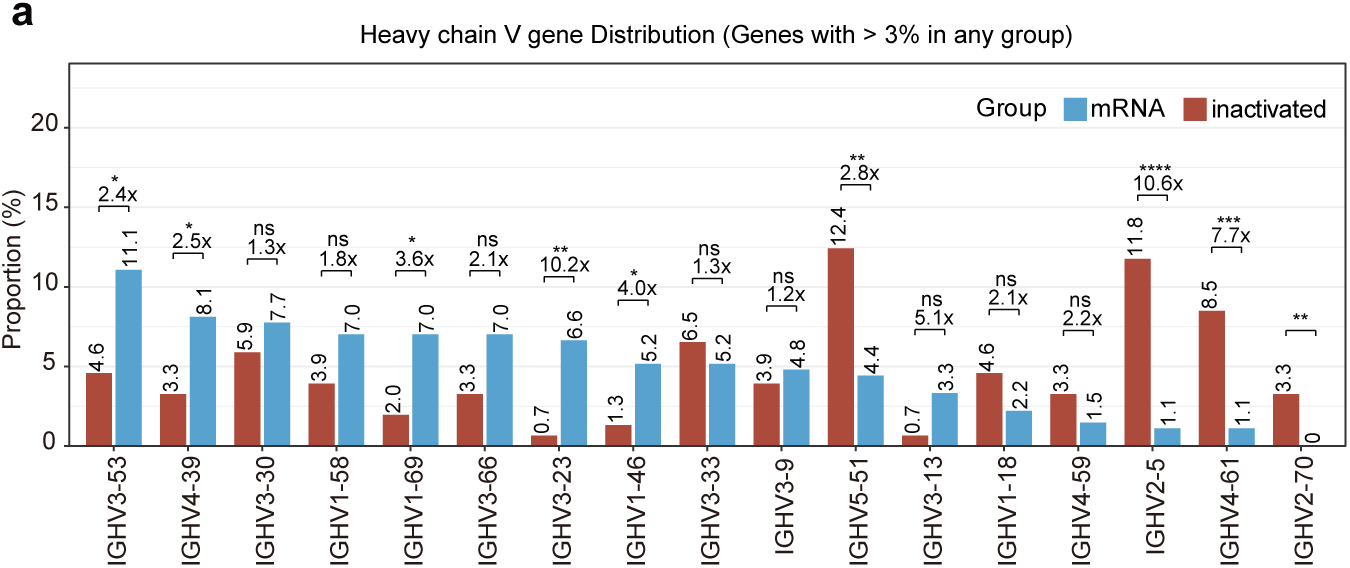
IGHV gene usage distinguishes highly- and weakly- imprinted antibody repertoires a, Frequency of heavy chain V gene usage of the two cohorts. Chi-square tests were used. \**P* < 0.05, \*\**P* < 0.01, \*\*\**P* < 0.001, \*\*\*\**P* < 0.0001; ns, not significant (P > 0.05).

**Extended Data Figure 5.**
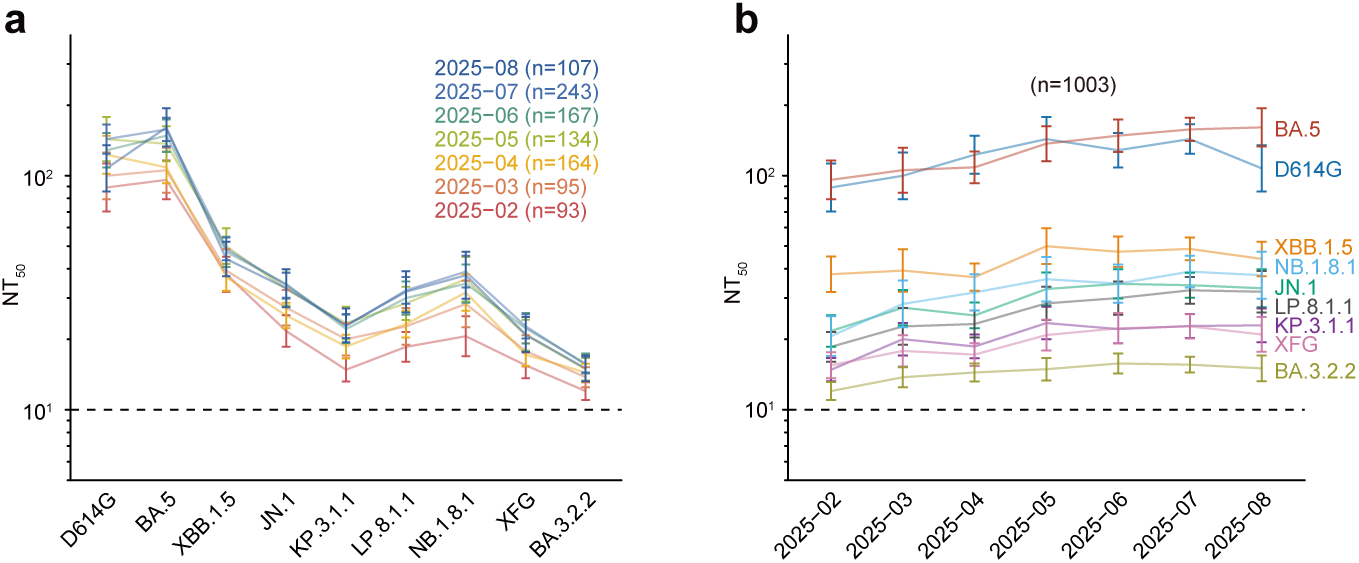
**Temporal increases in plasma neutralisation responses across the sampled population a-b**, Temporal changes in sampling time and plasma neutralising antibody responses against a panel of SARS-CoV-2 variant pseudoviruses. Geometric mean titres (GMTs) are shown on the top. Dashed lines indicate the limit of detection (NT_50_ = 10). Data are presented as geometric mean titres (GMT), with error bars indicating geometric standard deviation.

**Extended Data Figure 6.**
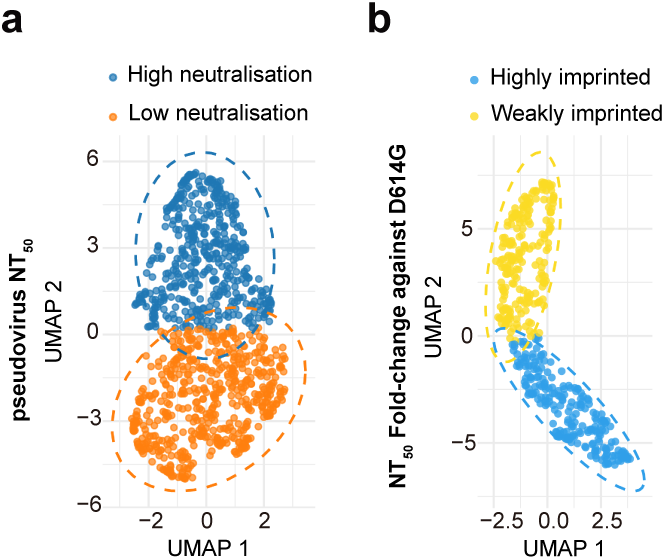
**Neutralisation-based UMAP clustering identifies imprinting-associated population groups a-b**, Uniform manifold approximation and projection (UMAP) projection showing clusters based on pseudovirus NT_50_ (**a**) and NT_50_ fold changes against D614G (**b**).

**Extended Data Figure 7.**
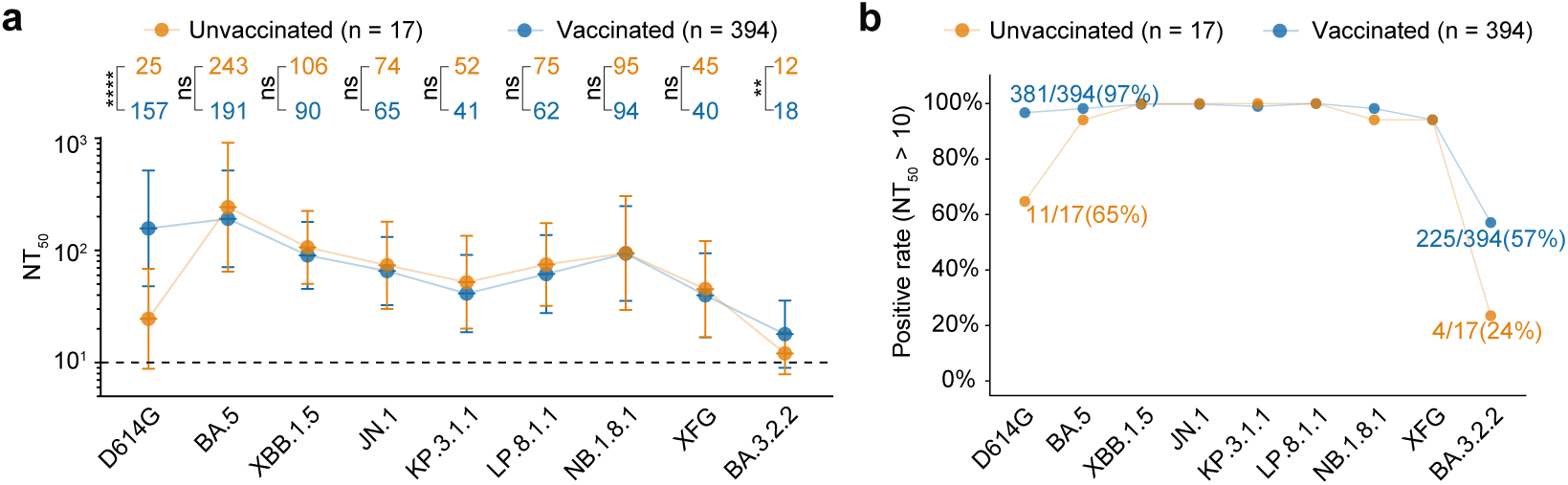
Unvaccinated individuals retain Omicron neutralisation but show reduced BA.3.2.2 activity. **a**, NAb response of the unvaccinated and vaccinated individuals against a panel of SARS-CoV-2 variant pseudoviruses. Geometric mean titres (GMTs) are shown on the top. Dashed lines indicate the limit of detection (NT_50_ = 10). Data are presented as geometric mean titres (GMT), with error bars indicating geometric standard deviation. **b**, Plasma neutralisation positivity rates for the data shown in **a**. Positivity was defined as a titre above the limit of detection (NT_50_ = 10). Two-tailed Wilcoxon rank-sum tests were used in **a**. \**P* < 0.05, \*\**P* < 0.01, \*\*\**P* < 0.001, \*\*\*\**P* < 0.0001; ns, not significant (P > 0.05).

**Extended Data Figure 8.**
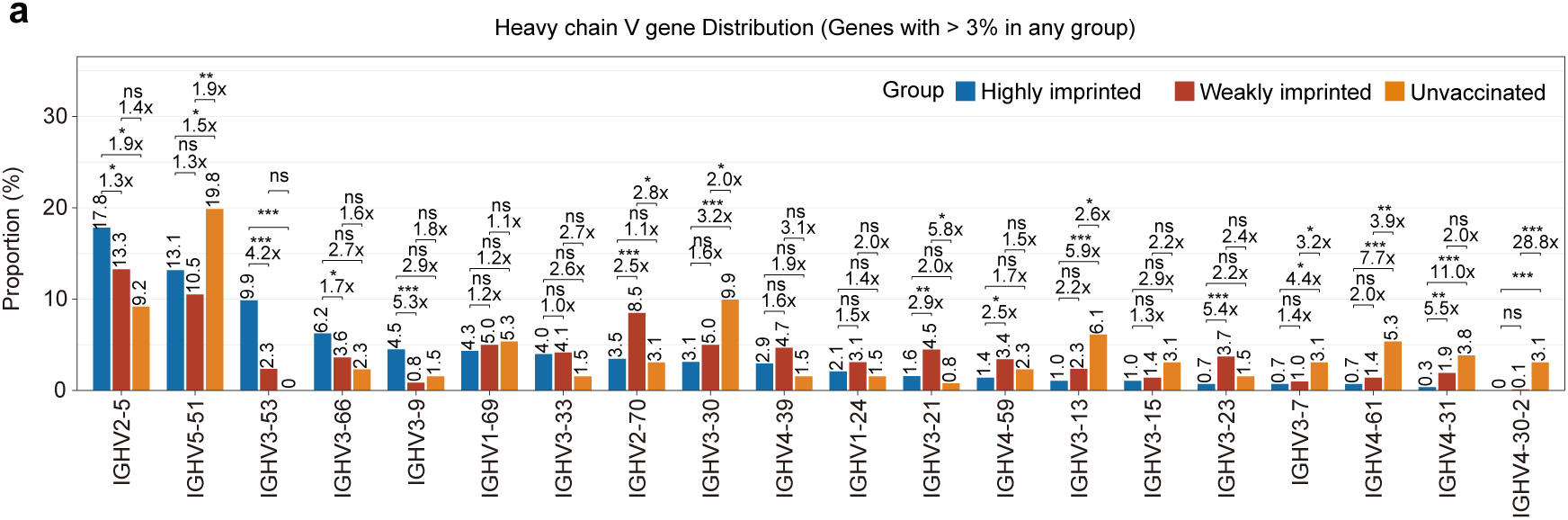
Expanded population BCR profiling supports imprinting-dependent IGHV usage a, Frequency of heavy-chain V gene usage in the highly imprinted, weakly imprinted and unvaccinated cohorts. Chi-square tests were used. \**P* < 0.05, \*\**P* < 0.01, \*\*\**P* < 0.001, \*\*\*\**P* < 0.0001; ns, not significant (P > 0.05).

